# Multi-state Ensemble Refinement for Occupancy Statistics (MEROS) in Time-Resolved X-ray Crystallography

**DOI:** 10.64898/2026.05.04.722701

**Authors:** Andreas Prester, Maria Spiliopoulou, Eike C. Schulz

## Abstract

Accurate determination of state occupancies is essential for interpreting the structural heterogeneity inherent in time-resolved crystallography. However, in cases of high spatial overlap between states, as commonly observed in time-resolved crystallography data, the strong correlation between occupancy and atomic displacement parameters (ADPs) can render single point estimates from standard refinement protocols unreliable. We introduce MEROS (Multi-state Ensemble Refinement for Occupancy Statistics), a pipeline that implements an ensemble refinement approach to assess the post-refinement occupancy–ADP statistics of multiple overlapping states. MEROS utilizes a Monte Carlo sampling of the parameter space, performing independent refinements from randomized starting occupancies and ADP values to empirically characterize the convergence and uncertainty of the solution. The method is implemented as a modular Python pipeline that wraps established refinement programs, ensuring compatibility with existing workflows. We demonstrate its applicability in two case studies: a two-state ligand binding model in T4 lysozyme L99A and a four-state covalent catalysis mechanism in *β*-lactamase CTX-M-14. MEROS provides occupancy and ADP mean values with standard deviations that directly quantify the informational content of the experimental diffraction data.

## Introduction

Time-resolved crystallography (TRX) has emerged as a powerful approach for visualising biochemical reactions at atomic resolution [1–3]. By initiating a reaction within protein crystals via photoactivation, rapid mixing, or temperature perturbation and collecting diffraction data at defined time delays, it is possible to structurally characterise transient reaction intermediates that are otherwise inaccessible or have previously only been observed in mutant variants [4– 7]. Serial crystallography approaches at synchrotrons (TR-SSX) and X-ray free-electron lasers (TR-SFX) have expanded the scope of time-resolved studies to a broad range of enzyme systems by reducing or eliminating radiation damage and by enabling data collection at physiologically relevant temperatures and timescales [8–12]. However, opposed to single discrete states TRX data fundamentally produces maps of overlapping structural ensembles, as each dataset represents a population-weighted superposition of multiple conformations within the crystal lattice. At any specific time point, the crystal comprises a mixture of unperturbed starting state, intermediate states, and perturbed state, each possessing a fractional occupancy that influence the observation of in-crystal reaction kinetics [13, 14]. Resolving this structural heterogeneity in dynamic crystallography therefore plays a central role for mechanistic interpretations [15, 16].

Multi-state structural modeling has been shown as an essential tool for interpreting electron density in perturbed datasets [17]. Central to this approach is the accurate representation of superpositions between ligand-bound and unbound states, a common feature of heterogeneous experimental datasets. Modeling the entire structural ensemble, rather than focusing on the perturbed state of interest, is critical for the high-quality refinement of low-occupancy states and is necessary to provide a comprehensive explanation of the observed density [17]. However, the calculation of occupancy values is complicated by high correlation of occupancy and atomic displacement parameters (*B* -factors), as both, low population and high flexibility, produce similar reductions in electron density. Standard refinement protocols often provide single point estimates for occupancy and *B*-factors that are sensitive to initial conditions, that may additionally contribute to subjectivity. Several computational approaches exist for analysing partially occupied states. Including PanDDA for detecting partial-occupancy ligand binding events [18], qFit for automated multi-conformer modelling [19], Xtrapol8 for structure-factor extrapolation to full occupancy [20], and singular-value decomposition methods for kinetic analysis of difference maps [21]. While these tools facilitate the modeling and the interpretation of TRX-data, neither provides a framework for quantifying the uncertainty of refined occupancy values in time-resolved multi-state models. To overcome this, a systematic sampling of the parameter space is required. By performing an ensemble of refinements from randomized starting values, the stability of the solution can be evaluated, allowing for the quantification of occupancy uncertainty and the identification of potential refinement bias.

The reliable determination of state occupancies is complicated by a fundamental parameter correlation in crystallographic refinement. Occupancy and *B*-factors (also atomic displacement parameters; ADPs) are intrinsically coupled, that is that reducing the occupancy of a group of atoms while simultaneously sharpening their displacement parameters can produce nearly identical contributions to the calculated structure factor amplitudes [22]. Consequently, changes in the occupancy can be partially offset by opposite changes in the *B*-factors, particularly at medium resolution, which is insufficient to distinguish between the two contributions. The problem is particularly acute in multi-state models where atoms from different states spatially overlap, as commonly occurs in enzyme active sites where successive catalytic intermediates share most of their atomic framework but differ in key chemical groups [23]. The practical consequence is that occupancy values reported in time-resolved crystallographic studies almost invariably lack uncertainty estimates. If these point estimates are used for subsequent analyses, such as fitting in-crystal kinetics or deriving reaction mechanisms, the resulting parameters exhibit a non-quantified systematic uncertainty, which undermines the statistical validity of the analysis.

To advance the rigorous analysis of time-resolved crystallography, we introduce MEROS (Multi-state Ensemble Refinement for Occupancy Statistics), a robust and accessible framework designed to extract occupancy values with quantitative uncertainty estimates from multi-state models. MEROS utilizes a Monte Carlo strategy to empirically assess the sensitivity of refined occupancies and *B*-factors to initial conditions. It adapts the ensemble-refinement verification approach demonstrated for simpler multi-state systems to the complex, overlapping-state sys-tems with time-resolved data [24]. The spread of the resulting distributions directly quantifies the extent to which the experimental data defines the population of each state. The method is designed to characterize structural heterogeneity within localized regions of interest, such as an enzyme active site or a hydrophobic cavity. Within this defined scope, the user assigns interconnected alternative conformations to functional groups or chemical states, for which occupancy values with error estimates are then obtained. To establish the scope and broad applicability of this method, we validated the pipeline on *in-situ* mixing serial crystallography data from two enzyme systems using the liquid-application-method for time-resolved crystallography (LAMA) spanning a range of multi-state complexity. Specifically, a two-state ligand binding event in T4 lysozyme L99A and a four state catalysis with a covalent intermediate in CTX-M-14 *β*-lactamase. MEROS is implemented as a Python pipeline and can be accessed at https://github.com/AndiPi-1/MEROS.

## MEROS

In standard crystallographic refinement, the target function, typically a maximum-likelihood function of the observed and calculated structure factor amplitudes, is minimised with respect to all model parameters, including atomic coordinates, occupancies, and *B*-factors. This yields point estimates at the minimum of the ML-function, but provides no direct information about the posterior probability distribution of these parameters given the data. MEROS takes an empirical approach by systematically probing the refinement landscape from a large number of diverse starting configurations. The hypothesis is that, if the data strongly constrain a particular state’s occupancy, independent refinements from all starting points converge to the same refined value, yielding a narrow distribution. Conversely, if the data are insufficient to fully resolve the occupancy–*B*-factor correlation, different starting configurations converge to a range of values, and the width of this distribution directly quantifies the resulting uncertainty. This approach treats the refinement process as an external optimization step, ensuring compatibility with various software programs while making minimal assumptions about the occupancy–*B*-factor distribution.

### Multi-state model definition

The multi-state model definition is crucial for the occupancy refinement as performed within the MEROS workflow. Using strong prior knowledge of specific functional states or intermediates provides a substantial analytical advantage. Stable intermediates that have been independently trapped and characterized in cryo-MX can be directly utilized to constrain and guide the interpretation of mixed-state electron density maps. This strategy applies equally to the initial reference states of the reaction cycle, defining the dark state in light-driven systems or the ligand-free apo state in diffusion-based experiments, as exemplified by T4L-L99A below. For optimal geometries, the reference models should ideally be derived from high- or ultra-high-resolution datasets. These are advantageous, as they define precise atomic coordinates, capture alternate conformations, and may even resolve individual protonation states. For each enzyme system, the initial multi-state model was constructed by placing the known state conformations (e.g. apo, substrate-bound, covalent intermediate, product) into the model. Each state has a defined ligand residue with an alternative conformation and, where necessary, associated protein conformational changes were modeled as alternate conformers. This ensures that the state specific residues/ligands do not interfere with those from another state.

For the *first model system*, T4L-L99A a shift in the *F*-helix residues to an open state has been observed, upon binding of indole in the hydrophobic cavity. We have therefore defined 2 states for the enzyme, the apo state with an empty cavity and the closed *F*-helix position, and the bound state with indole in the hydrophobic cavity as well as the *F*-helix in the open state (**Figure 5 A**). This makes three occupancy groups. Groups one and two consist of the *F*-helix residues (107–115) in alternative conformation A and B, respectively. During refinement these are constrained to sum to 100 % occupancy as they are enzyme residues. The third occupancy group consists of the bound ligand indole, which is not constrained to 100 %.

When the *second model system*, CTX-M-14 is mixed with piperacillin, we can build 4 different states in the active site of the enzyme (**Figure 6 A**). Modeling the native state represented by water molecules, and in this case a sulfate, in the active site can be crucial for the correct interpretation and refinement of the data, especially when ligand occupancy is low [17]. The second state is a Michaelis-Menten complex with the unbound substrate piperacillin in the active site. The ligand has displaced the water molecules and the sulfate of the native site. Information about this state was obtained from a S70G mutant variant. In the third state the Ser70 side chain hydroxyl group has performed a nucleophilic attack on the carbonyl-carbon atom of the piperacillin *β*-lactam ring and formed a covalent acyl-enzyme intermediate as obtained from E166A mutant variants. The fourth state is formed after hydrolysis via the catalytic water molecule and resembles the product state complex with a formed carboxylate group at the piperacillin and a rotated Ser70. Since the last two states involve alternative conformations of the Ser70, we have modeled 4 alternative conformers of Ser70 and assigned one to each state. MEROS then operates in these multi-state models and quantifies how well the experimental data fit each states occupancy. Each states’ reference structure was obtained from crystals grown with the same strategy and conditions as for the time-resolved experiment.

### The MEROS algorithm

The MEROS procedure consists of three steps (**Figure 1**):

**Figure 1:**
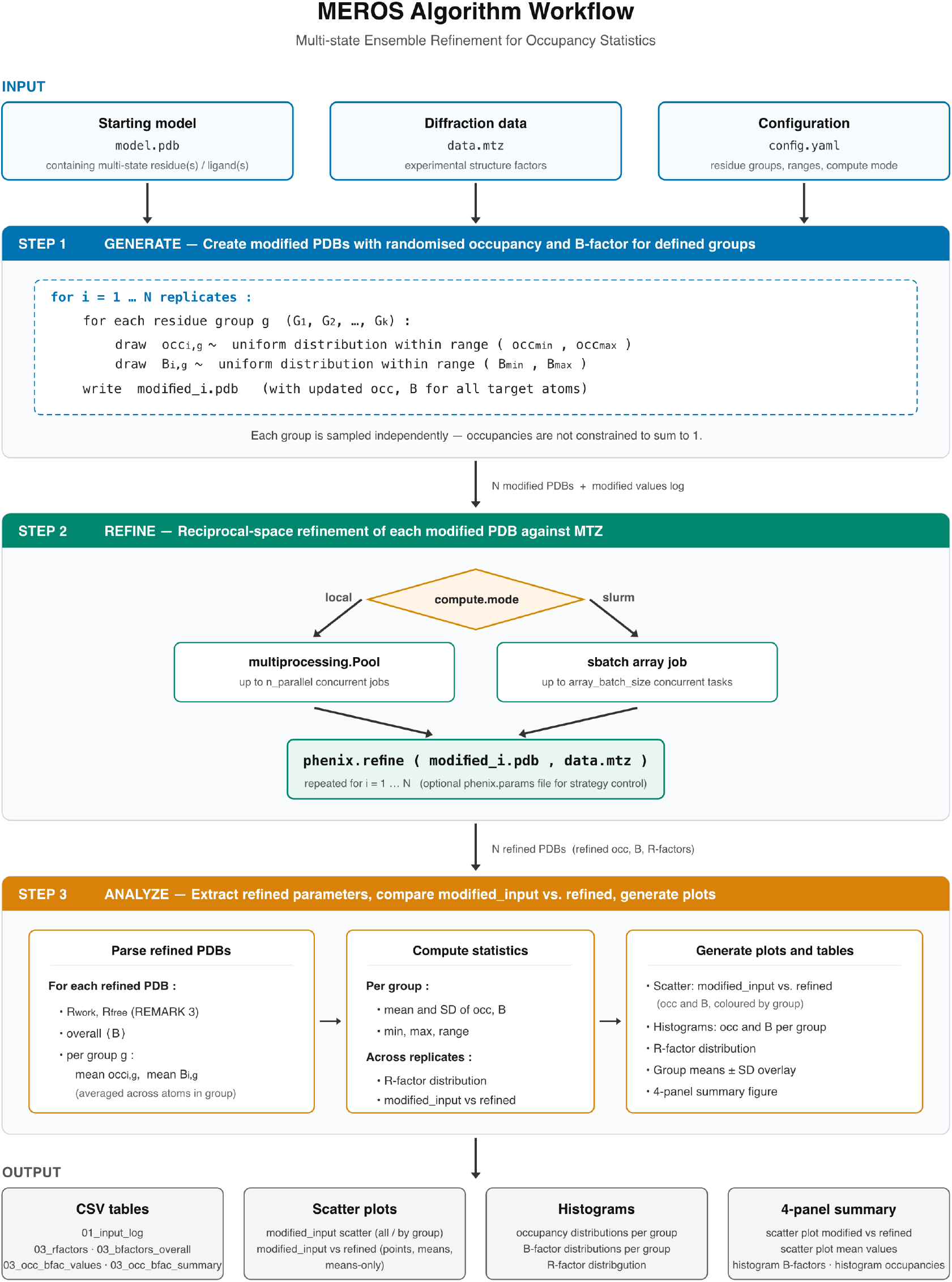
Overview of the MEROS computational workflow applied for time-resolved crystallography.

#### Step 1: Ensemble generation

Given a multi-state model *ℳ*_0_ with *K* user-defined residue groups, MEROS generates *N* modified copies. For each copy *i* ∈ {1,. .., *N*} and each group *k* ∈ {1,. .., *K*}, occupancy and *B*-factor values are drawn independently from uniform distributions:

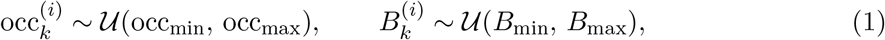

where the occupancy range [occ_min_, occ_max_] and *B*-factor range [*B*_min_, *B*_max_] are user-configurable (defaults: [0.01, 0.99] and [10, 50] Å^2^, respectively). The drawn values replace the occupancy and isotropic *B*-factor of every atom in group *k* in the PDB coordinate file, while all other atoms remain unchanged. Each modified model receives a unique (occ_*k*_, *B*_*k*_) combination per group to ensure comprehensive coverage of the parameter space. The uniform prior over the full occupancy range ensures that MEROS explores regions of parameter space that might not be considered in conventional refinement, avoiding confirmation bias. The scatter plots provide visual validation that the generated values are evenly distributed within the given range (**Figure 2**).

**Figure 2:**
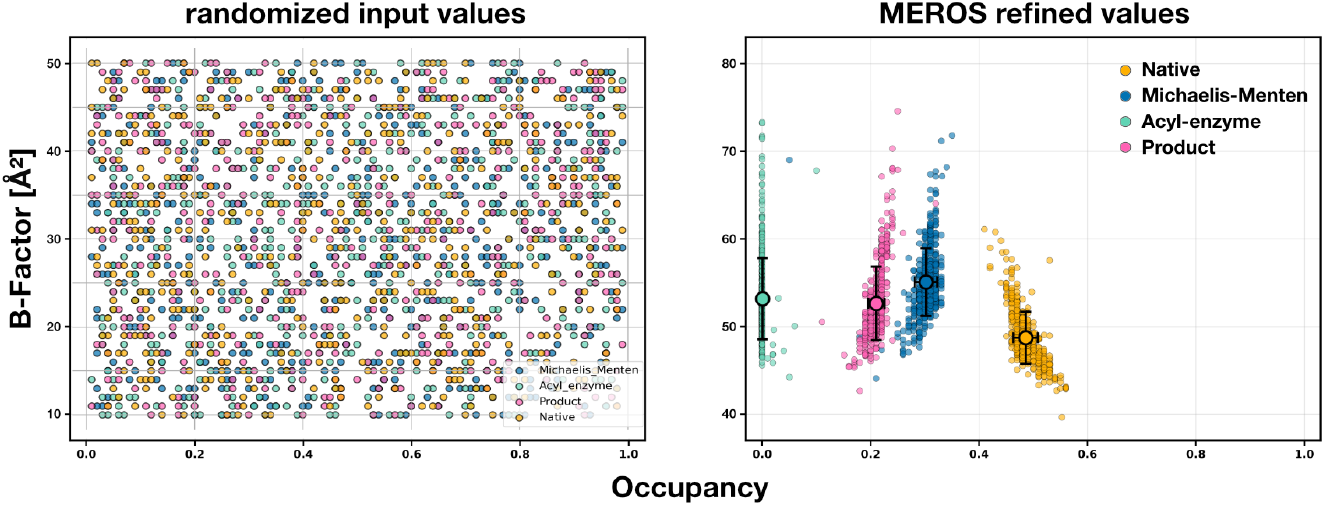
Occupancy–*B*-factor distributions of the modified random input values and the MEROS refined values. Scatter plots show the randomized input values for each defined group/state with an even distribution withing the given range. Following MEROS refinement, the occupancy versus *B*-factor distribution for the CTX-M-14 catalysis steps, including the apo state (yellow), the Michaelis–Menten state (blue), the acyl–enzyme intermediate state (teal), and the hydrolysed product state (pink), cluster around their refined values. Each point represents one of 500 independent refinement runs with the randomized starting parameters. Distinct clusters highlight the refinement behavior and occupancy convergence of each state, reflecting the stages of enzymatic catalysis.

#### Step 2: Independent batch refinement

Each of the *N* modified models is refined against the experimental reflection data (MTZ) using the same refinement strategy, restraints, and parameter settings as would be applied in a standard refinement. Currently, all refinements were performed with phenix.refine [25] using identical parameter files for all replicates within a dataset. Importantly, occupancy, *B*-factors, and atomic coordinates are all free to refine; none is held fixed. However, the refinement is performed using shared occupancy constraints for each residue group to reflect the physical presence of discrete structural states. By allowing individual *B*-factors for every atom within a group, the model could accommodate local flexibility without compromising the states occupancy estimate, thereby mitigating potential biases caused by non-uniform mobility across the group. This ensures that each modified model finds its own local optimum in the full parameter space. Refinements are executed in parallel using either local multiprocessing or SLURM cluster array jobs, enabling high-throughput processing of hundreds of replicates within hours on modern hardware.

#### Step 3: Statistical analysis

After refinement, MEROS extracts the converged occupancy and *B*-factor values for every residue group from each refined PDB. For groups comprising multiple atoms, the group *B* - factor is taken as the arithmetic mean over all constituent atoms. Additionally, *R*_work_ and *R*_free_ values, and overall mean *B*-factors are extracted from the refinement output PDB headers as a control for failed refinements. For each group *k*, the following summary statistics are computed from the sample of size *N* :

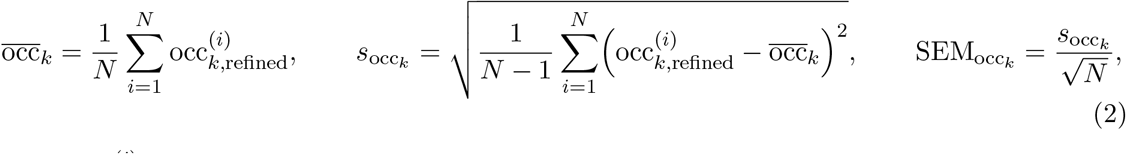

where 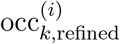 denotes the refined occupancy of group *k* in replicate *i*. The values for the *B*-factors were determined accordingly. The distributions are visualized as histograms and scatter plots (**Figure 2 and 3**).

**Figure 3:**
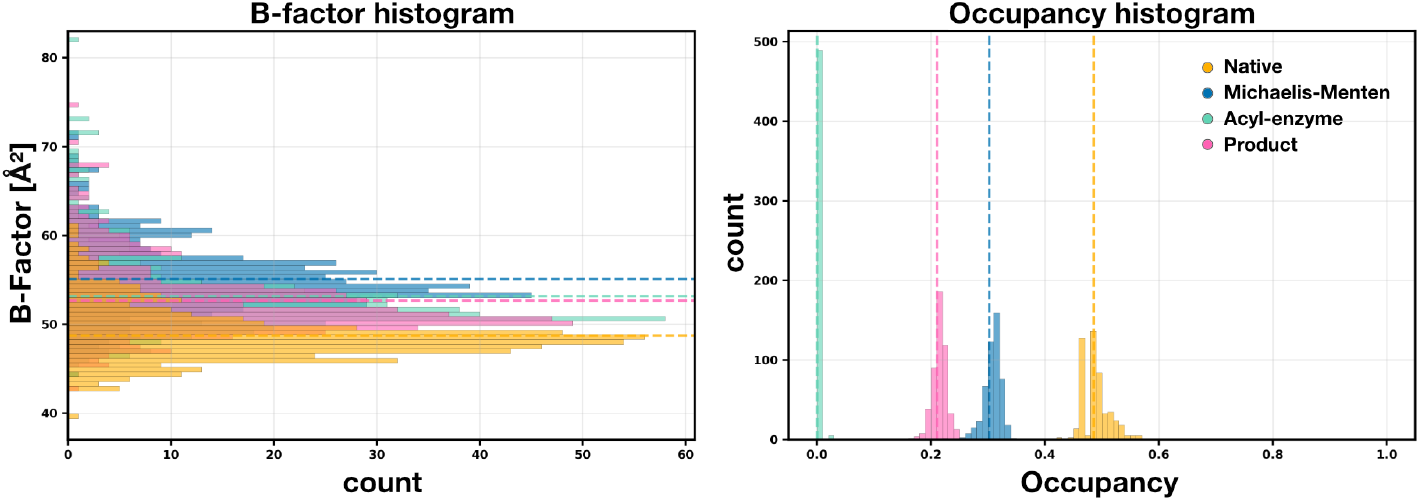
Histograms showing the MEROS refined value distributions for Occupancies and *B*-factors. Scatter plots can appear cluttered at high replicate numbers, which can create the impression of a noisy distribution by highlighting the outliers and obscuring multiple overlapping points around the mean. For this reason, separate histograms are generated for the occupancy and *B*-factor distributions, in order to better visualise the distribution as a function of the counts. The mean values for each group and each distribution are shown as a dashed lines. The CTX-M-14 catalysis steps are defined as the apo state (yellow), the Michaelis–Menten state (blue), the acyl–enzyme intermediate state (teal), and the hydrolysed product state (pink).

### Implementation and workflow

MEROS is implemented as a modular Python pipeline consisting of the following components:

- `meros.py:` command-line entry point with argument parsing and pipeline orchestration;
- `config.py:` YAML configuration loading with validation and default-setting;
- `generate.py:` generation of modified PDB files with randomised occupancy and *B*-factor values for defined residue groups;
- `refine.py:` refinement execution via local multiprocessing or SLURM cluster array jobs using established refinement programs;
- `analyze.py:` extraction of refined parameters, statistical analysis, and visualisation;
- `pdb utils.py:` shared PDB file parsing utilities.

The pipeline is configured via a single YAML file that specifies input files, residue group definitions, generation parameters, refinement strategy, compute resources, and plot settings. A multi-dataset mode allows processing of multiple time points (or experimental conditions) in a single configuration file, applying the same group definitions and analysis parameters to ensure consistency across the time series. MEROS wraps the refinement program as a black box, passing each modified PDB together with the experimental MTZ and a user-supplied parameter file. This architecture ensures compatibility with established refinement programs without re-implementing the likelihood function. Currently, `phenix.refine` [25] is supported as the primary refinement program. However, due to its modular design MEROS can also accommodate an extension to other programs (e.g. `REFMAC5` [22] or `BUSTER` [26]). The software requires Python *↗* 3.8 with NumPy, pandas, Matplotlib, and PyYAML.

## Results

To validate the MEROS approach, we first examined the general behaviour of the occupancy– *B*-factor refinement landscape. The key diagnostic output of MEROS is the two-dimensional scatter plot of refined occupancy vs. *B*-factor for each residue group (**Figure 2**). When the experimental data strongly constrain the occupancy of a state, all *N* refinements from diverse starting points converge to essentially the same refined values, producing a tight cluster. When the occupancy and *B*-factor are poorly resolved, due to low occupancy, limited resolution, poor data quality, or too much spatial overlap between states, the converged values form an elongated distribution along the occupancy–*B*-factor, directly visualising the parameter covariance. As an internal consistency check, MEROS monitors *R*_work_ and *R*_free_ across all replicates. A narrow, consistent distribution of *R*-factors confirms that all replicates reach comparable fit quality and that differences in refined occupancies do not reflect failed or trapped refinements but genuine parameter ambiguity (**Figure 4**).

**Figure 4:**
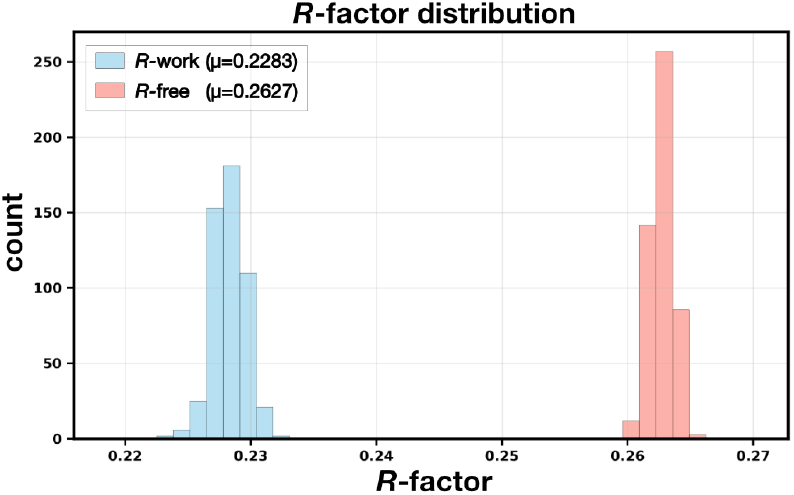
*R*-factor distribution of MEROS refinement of CTX-M-14 mixed with piperacillin at 20 °C, after a delay time of 3 seconds, with *N* = 500 refinements. The histogram shows no significant outliers, indicating a generally stable refinement.

**Figure 5:**
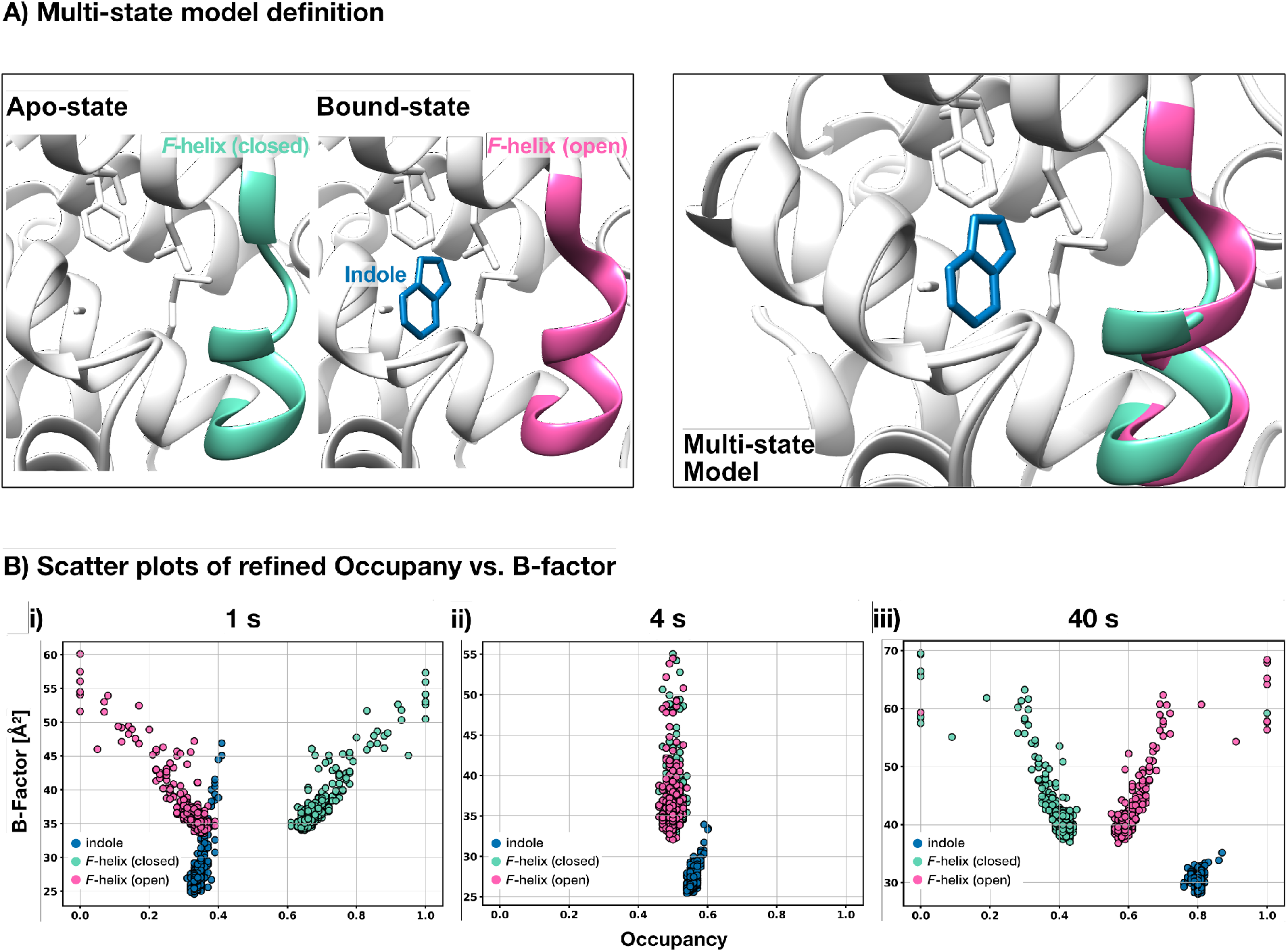
**A)** Group definition for T4L-L99A with indole of the MEROS workflow in time-resolved crystallography. **B)** Occupancy–*B*-factor distributions from independent refinements. Scatter plots of refined occupancies versus *B*-factors for indole (blue) and the *F*-helix in closed (teal) and open (pink) conformations for certain time points: 1 s **(i)**, 4 s **(ii)**, and 40 s **(iii).** Each point represents one of 500 independent refinement runs with randomized starting parameters. Distinct clusters highlight the refinement behavior and occupancy convergence of each group in consistence with increasing indole and open *F*-helix conformations, the occupancy of the closed *F*-helix conformation decreases over time.

**Figure 6:**
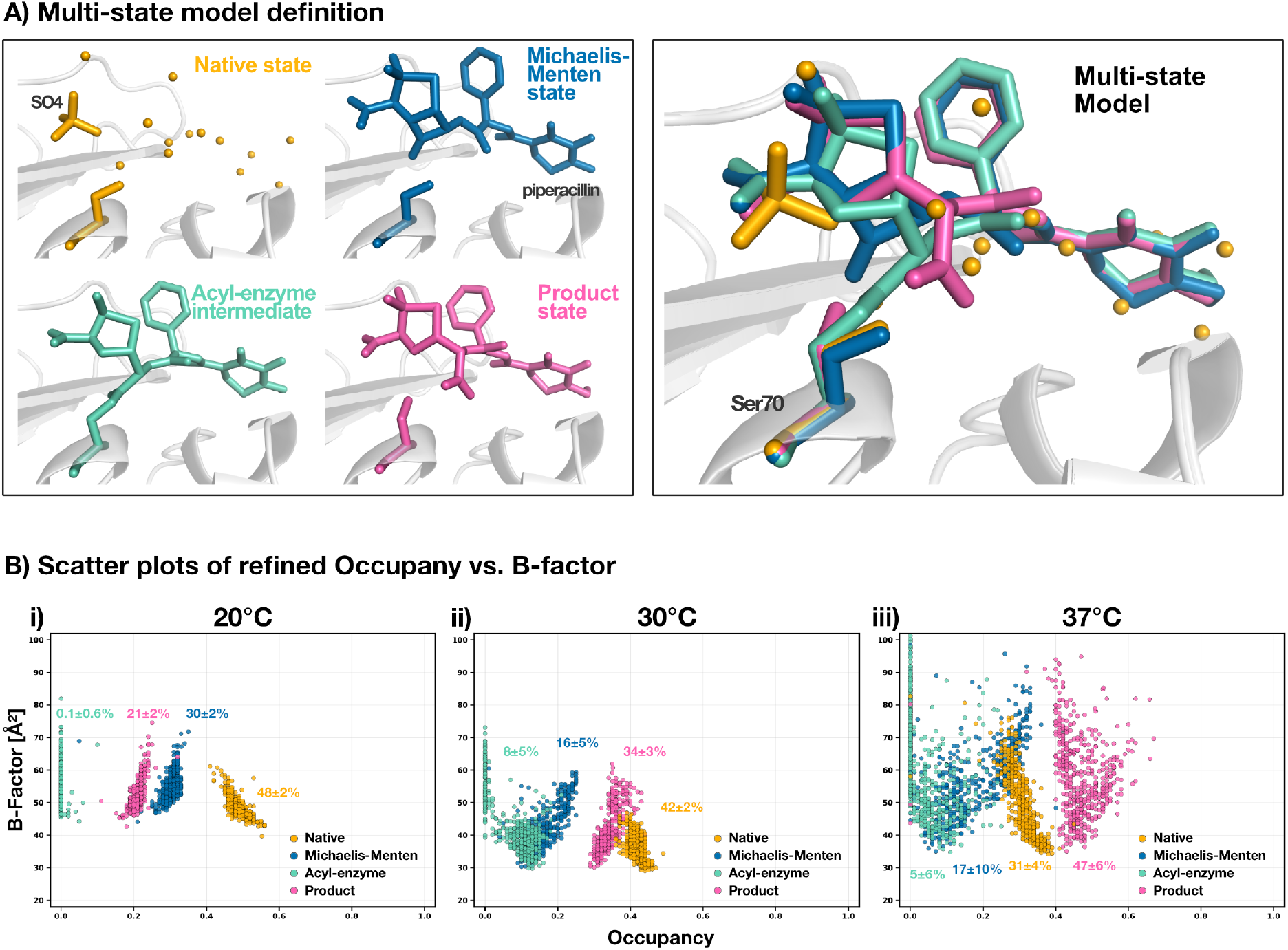
**A)** Group definition for CTX-M-14 piperacillin of the MEROS workflow in time-resolved crystallography. **B)** Occupancy–*B*-factor distributions from independent refinements showed via scatter plots of refined occupancies versus *B*-factors for CTX-M-14 catalysis steps. The apo state (yellow), the Michaelis-Menten state (blue), the Acyl-enzyme intermediate state (teal) and the hydrolysed product state (pink) conformations are shown for time point 3 s at various temperatures: 20 °C **(i)**, 30 °C **(ii)**, and 37 °C **(iii).** Each point represents one of 500 independent refinement runs with randomized starting parameters. Distinct clusters highlight the refinement behavior and occupancy convergence of each state, reflecting the influence of temperature on the enzymatic catalysis.

### T4 lysozyme: validation on a two-state binding system

The T4 lysozyme L99A mutant provides an ideal validation system because it presents a simple two-state scenario: the hydrophobic cavity at position 99 is either empty (apo state) or occupied by a small-molecule ligand (bound state) [27, 28]. Here we use our TR-SSX data on T4L-L99A mixed with indole using the LAMA method at 3 delay times (1 s, 4 s, 40 s) [29]. At saturating ligand concentrations, the occupancy approaches unity, providing a known endpoint for validation.

The two-state model comprises three occupancy groups, the *F*-helix residues (107–115) in the closed and open conformations, which are constrained to sum to unity as they are enzyme residues. The bound indole ligand, is refined independently without summing to unity.

At all three delay times, all three groups produce compact, well-defined clusters in the occupancy–*B*-factor scatter plots (**Figure 5 B**). This is consistent with the relatively simple geometry of the T4L system, in which the ligand occupies a previously empty pocket. As a result, the data constrain the occupancy–*B*-factor correlation more tightly, and MEROS resolves well-defined distributions from all *N* = 500 starting configurations. Notably, the *F*-helix closed cluster starts as the highest occupied group and progressively shifts to lower occupancy. Its distribution broadens slightly at the 40 s time point, where the population of this state is small.

Across the three delay times, the indole occupancy increases from 34(±1) % at 1 s to 56(±1) % at 4 s and 80(±1) % at 40 s, approaching saturation at the longest delay. The open *F*-helix occupancy follows the same trend, rising from 32(±6) % at 1 s to 50(±1) % at 4 s and 60(±6) % at 40 s, and the closed *F*-helix occupancy decreases correspondingly from 68(±6) % to 50(±1) % and 40(±6) %, satisfying the sum-to-unity constraint in refinement at all time points. At the earlier time points the indole and the open *F*-helix occupancies are in close agreement within their respective uncertainties, consistent with the expected coupling between ligand binding and *F*-helix opening. However, at 40 s the indole occupancy (80(±1) %) exceeds the open *F*-helix occupancy (60(±6) %) by approximately 20 percentage points, and the uncertainty on the *F*-helix groups is notably larger than that on indole. This difference in precision is consistent with the differing degrees of spatial overlap in the model. Indole occupies a previously empty cavity with no competing group, so its electron density is uniquely attributable to the bound state and the refinement converges tightly. In contrast, the open and closed *F*-helix conformations occupy overlapping regions of the protein and compete for the similar electron density. This weakens the data constraint on their individual occupancies and broadens the MEROS distributions. Furthermore, the *F*-helix group (residues 107–115) comprises significantly more atoms than the small indole molecule. This introduces a greater number of *B*-factors, which collectively contribute to the parameter correlation and further increase the uncertainty relative to the single-ligand group.

The key contribution of MEROS is the standard deviation on each calculated mean value, which provides a quantitative measure of how tightly the experimental data define the indole and *F*-helix occupancies at each time point. The narrow distributions at all three delay times confirm that the T4L system is well-suited for validation purposes, and that the MEROS-reported uncertainties reflect parameter precision. This information is inaccessible from a conventional single refinement. Furthermore, by applying MEROS across a larger series of delay times, the occupancy values can be assembled into an in-crystal binding curve. This enables a direct and quantitative description of ligand uptake kinetics within the crystal lattice.

#### CTX-M-14: four-state covalent catalysis of piperacillin

CTX-M-14 is a class A *β*-lactamase that hydrolyses *β*-lactam antibiotics through a mechanism involving formation (acylation) and breakdown (deacylation) of a covalent acyl-enzyme intermediate [30]. In time-resolved experiments with piperacillin as the substrate, the crystal contains four coexisting states: (1) the native (substrate-free) active site (2) the non-covalent Michaelis– Menten complex, (3) the covalent acyl-enzyme intermediate at Ser70, and (4) the hydrolysed product [12]. This is a more demanding test case, involving four partially overlapping states, the combined occupancies of which should not exceed one. Here we show a temperature series of CTX-M-14 mixed with piperacillin at a delay time of 3 seconds at 20 °C, 30 °C, and 37 °C, we described previously [12].

The four chemical states occupy overlapping regions of the active site and are distinguished by a combination of ligand chemistry and Ser70 side-chain geometry (**Figure 6 A**). In the Michaelis–Menten complex, piperacillin is bound non-covalently and Ser70 retains its ground-state rotamer. Upon nucleophilic attack, the acyl-enzyme intermediate forms a covalent ester bond between the Ser70 O*ε* and the piperacillin carbonyl carbon, which requires Ser70 to adopt a distinct rotamer rotated towards the oxyanion hole. In the product state, the covalent bond is hydrolyzed and Ser70 turns further to a distinct post-deacylation rotamer. Additionally, hydrolysis converts the acyl carbonyl to a carboxylate, which adopts a different position in the active site, providing a structural handle for discriminating the product from the acylenzyme intermediate in the electron density. Because MEROS refines each of the four states independently from randomised starting parameters, the resulting scatter plots directly reveal both the quality of convergence and the correlation of the occupancy and *B*-factor for each state.

At all three temperatures the native and product states produce well-defined, compact clusters in the occupancy–*B*-factor scatter plots (**Figure 6 B**). This reflects the comparatively high electron-density signal from these states. The reactive intermediates, Michaelis–Menten complex and acyl-enzyme, exhibit progressively more elongated distributions with increasing temperature, consistent with a reduction in occupancy and a concomitant weakening of the data constraint. At 20 °C the acyl-enzyme cluster collapses to near-zero occupancy with a very narrow distribution, indicating that the data exclude any significant population of the covalent intermediate in this dataset at this temperature. At 30 °C and 37 °C the cluster broadens along the occupancy axis, reflecting a genuine ambiguity that MEROS resolves into quantified uncertainty rather than an unverified point estimate.

MEROS reports occupancies with standard deviations from *N* = 500 independent refinements. The calculated values for all four states at each temperature are listed in **Table 1**. With the four-state constrained occupancy model, the sum of all state populations is maintained at unity, and in all cases the MEROS means satisfy this constraint within rounding errors. At 20 °C the native active site is the dominant species at 49(±2) %, the Michaelis–Menten complex accounts for 30(±2) %, the hydrolysed product for 21(±2) %, and the acyl-enzyme intermediate is essentially absent at 0.1(±0.6) %. The low acyl-enzyme occupancy at 20 °C suggests that acylation may be the rate-limiting step at this temperature. Under this interpretation, the 21% product already present after 3 s at 20 °C indicates that deacylation has occurred. This im-plies that the acyl-enzyme intermediate is consumed by deacylation faster than it accumulates, keeping its steady-state population low. However, a conclusive explanation of the reaction and identification of the rate-limiting step would require more data, as well as direct kinetic measurements. At 30 °C the acyl-enzyme intermediate becomes observable at 9(±5) %, while the Michaelis–Menten complex decreases to 16(±5) % and the product fraction rises to 34(±3) %. This pattern is consistent with an overall acceleration of the acylation–deacylation cycle relative to 20 °C. At 37 °C the occupancy distribution is most shifted towards the product state at the 3 s time point with 47(±6) %. The native states occupancy is at 31(±4) % and the acylenzyme intermediates at 5(±6) %. This value whose standard deviation is comparable to the estimate itself, underscoring the marginal signal-to-noise for this state under these conditions. The standard deviation of this value is comparable to the calculated mean itself, highlighting the marginal signal-to-noise ratio for this state under these conditions. The Michaelis–Menten complex at 37 °C yields an occupancy of 17(±10) %, the largest relative uncertainty in the dataset, which may reflect a more transient population.

**Table 1:** Comparison of 3 different starting points for a regular refinement and MEROS refinement of CTX-M-14 mixed with piperacillin at 3.0 s delay time. The reference output is basically the initial finalized structure that was then used as the starting structure for MEROS. The Output 25% is the same structure but with the occupancy set to 25 % and *B*-factors to 30 Å^2^ for each group, resulting in the same starting parameters for each group. The Output-mean-input is the result of a single refinement using the MEROS mean values as starting parameters for occupancy and *B*-factor.

| Temp | Native |  |  |  | Michaelis Menten |  |  |  | Acyl enzyme |  |  |  | Product |  |  |  |
| --- | --- | --- | --- | --- | --- | --- | --- | --- | --- | --- | --- | --- | --- | --- | --- | --- |
|  | Output reference | Output 25% | Output mean input | MEROS mean | Output reference | Output 25% | Output mean input | MEROS mean | Output reference | Output 25% | Output mean input | MEROS mean | Output reference | Output 25% | Output mean input | MEROS mean |
| *Occupancy (%)* |  |  |  |  |  |  |  |  |  |  |  |  |  |  |  |  |
| 20 °C | 54 | 49 | 48 | 49 ± 2 | 27 | 31 | 30 | 30 ± 2 | 0 | 0 | 0 | 0 ± 1 | 19 | 22 | 22 | 21 ± 2 |
| 30 °C | 45 | 45 | 46 | 42 ± 2 | 13 | 13 | 14 | 16 ± 5 | 11 | 11 | 10 | 9 ± 5 | 31 | 31 | 30 | 34 ± 3 |
| 37 °C | 38 | 36 | 38 | 31 ± 4 | 15 | 14 | 14 | 17 ± 10 | 6 | 5 | 3 | 5 ± 6 | 42 | 44 | 44 | 47 ± 6 |
| *B-factor (Å<sup>2</sup>)* |  |  |  |  |  |  |  |  |  |  |  |  |  |  |  |  |
| 20 °C | 45 | 49 | 48 | 49 ± 3 | 50 | 56 | 54 | 55 ± 4 | 49 | 53 | 52 | 53 ± 5 | 47 | 52 | 51 | 53 ± 4 |
| 30 °C | 30 | 29 | 29 | 36 ± 4 | 32 | 32 | 31 | 42 ± 7 | 32 | 31 | 30 | 43 ± 9 | 32 | 31 | 30 | 42 ± 7 |
| 37 °C | 36 | 36 | 35 | 50 ± 12 | 39 | 42 | 42 | 58 ± 12 | 39 | 41 | 41 | 60 ± 16 | 39 | 41 | 40 | 58 ± 13 |

The temperature-dependent increase of intermediate state occupancies at a fixed delay time is consistent with the acceleration of reaction rates at higher temperature [31]. The acyl-enzyme intermediate, though short-lived, is detectable at 30 °C and 37 °C, confirming that the experimental conditions capture a genuine kinetic snapshot of the covalent catalysis cycle in the crystal. Critically, MEROS not only reports these values but also exposes when the diffraction data are insufficient to robustly define a particular state. The large standard deviation on the acyl-enzyme occupancy at 37 °C is a direct indication that any subsequent analysis must propagate this uncertainty. Without MEROS, such an estimate would be presented as a precise, apparently trustworthy number derived from a single refinement. Together, the three temperature points illustrate that MEROS can track the progression of a multi-step enzymatic reaction across a temperature series, providing both the mean occupancy trajectory and the confidence with which each point on that trajectory is defined by the experimental data.

#### Comparison of conventional single refinement with MEROS refinement

To assess the influence of initial conditions on refinement outcomes and to benchmark MEROS directly against conventional single-structure refinement, we compared four approaches applied to the CTX-M-14 piperacillin data at a 3 s delay time at 20 °C, 30 °C, and 37 °C: (*i*) the *reference output*, representing the finalized multi-state model that also served as the starting structure for MEROS; (*ii*) a refinement initialized with identical uniform starting parameters for all groups (occupancy 25 % and *B*-factor 30 Å^2^); (*iii*) a single refinement using the MEROS ensemble means as starting parameters (*output mean input*); and (*iv*) the MEROS ensemble mean with standard deviation derived from *N* = 500 independent refinements. All four procedures used the same phenix.refine strategy and parameter file (**Table 1**).

The comparison reveals two principal findings concerning occupancy and *B*-factor refinement behaviour. For occupancy values, all three single-refinement approaches yield closely consistent results that agree well with the MEROS mean across all groups and temperatures. The three distinct starting conditions converge to within 2–3 percentage points of each other in virtually every case, and the largest discrepancy between a single-refinement estimate and the MEROS mean is approximately 7 percentage points (Native state at 37 °C: 38 % vs. 31(±4) %). This starting-point independence demonstrates that occupancy refinement is robust for this system, and that a conventional single refinement provides a reliable point estimate. In contrast, *B*-factor values exhibit a more pronounced sensitivity at elevated temperatures. At 20 °C, single-refinement and MEROS *B*-factors agree within 3–6 Å^2^ for all groups. At 30 °C and 37 °C, however, the MEROS means consistently exceed single-refinement values by up to 14 Å^2^, with substantially larger standard deviations. The most likely explanation for this temperature-dependent divergence is a partial convergence artefact within the MEROS ensemble rather than a systematic bias in the single-refinement estimates. MEROS draws starting occupancies and *B*-factors independently from broad uniform distributions and does not impose a constraint that occupancies sum to unity at the input stage. Thus, a minority of replicates will begin from highly unfavourable configurations. For example, with near-zero occupancy for a state that carries a substantial population, or with starting *B*-factors far above the true value. At 20 °C, where all four states are comparatively well-scattering and the data are informative, the refinement corrects even these poor starting points efficiently, and the multi-data mean matches the single-refinement result closely. At higher temperatures, where faster catalytic turnover can reduce the occupancy of individual, short-lived states and the signal-to-noise of the electron density decreases, a fraction of these challenging starting points may not fully converge to the global minimum within the allocated refinement cycles. Instead they settle in local minima characterised by elevated *B*-factors that partially compensate for the misguided occupancy input assignment. These sub-optimal replicates are captured directly in the broader *B*-factor distributions and the enlarged standard deviations observed at 30 °C and 37 °C, and are precisely what motivates inspection of the full scatter plot as well as the histograms as a convergence diagnostic. The shape of the resulting *B*-factor distributions is characteristically non-Gaussian, exhibiting a pronounced shoulder toward elevated *B* -values, as is clearly visible in the MEROS histograms (**Figure S1**). This asymmetry has a physical origin as atomic displacement parameters are restricted to positive values *B >* 0, and thermal motion establishes a practical minimum of several Å^2^ even in the most ordered atoms. Contrary to this, no equivalent upper bound exists, so the distribution cannot develop a symmetric shoulder at low *B* -values to balance the one observed at high *B* -values. This asymmetry is further amplified by the directional nature of the occupancy–*B*-factor correlation. When a replicate starts and converges to a solution with an overestimated occupancy for a given state, the refinement compensates by elevating the *B*-factor of that state to reduce its calculated scattering contribution back to the level supported by the data. The inverse, effect is not as pronounced, as it is limited by to the lower *B*-factor limit. The result is a primary distribution peak centered near the well-converged solution, with a pronounced shoulder extending toward elevated *B* -values that represents the sub-population of partially-converged replicates. Because the majority of replicates do converge correctly, the mean remains a valid estimate. However, the *B*-factor means at elevated temperature are pulled slightly upward relative to a fully converged single refinement. The most important contribution of MEROS to this comparison is nonetheless the quantification of uncertainty, which is entirely inaccessible from any single refinement regardless of the choice of starting conditions. The standard deviations reveal contrasting levels of parameter precision across states and temperatures. The Native and Product states are well-constrained at all three temperatures, with occupancy standard deviations of 2–6 %, consistent with their higher fractional populations. The reactive intermediates become progressively less defined with increasing temperature. The Michaelis–Menten complex at 37 °C reports an occupancy of 17(±10) %, and the acyl-enzyme intermediate at the same temperature reports 5(±6) %, a case where the uncertainty is comparable to the value itself. For *B*-factors, the acyl-enzyme intermediate at 37 °C reaches *θ* = 16 Å^2^, the largest uncertainty in the dataset, further reflecting the difficulty of resolving this low-occupancy state at elevated temperature. These results illustrate that MEROS provides consistent mean values and also indicates when the experimental data are insufficient to reliably distinguish between individual states. This information is crucial for to correct interpretation of temperature-dependent occupancy changes and for any downstream kinetic analysis.

## Discussion

The growing use of time-resolved serial crystallography has enabled the routine observation of dynamic enzyme-reactant interactions, using transient changes in electron density to map reaction pathways. However, as the field transitions from evaluating qualitative structural snapshots to conducting quantitative kinetic analyses, the ability to precisely quantify fractional occupancies becomes paramount. Because TRX datasets represent population-weighted superpositions of multiple states, robust uncertainty estimation is essential for meaningful interpretation.

The practical necessity of this quantification is best illustrated by the statistical distinction between reaction steps. For example, two point estimates of, for instance, *ôcc*_1_ = 0.35 and *ôcc*_2_ = 0.50 suggest a 15% progression of a reaction. However, if the associated standard deviations are *θ* = 0.16, these states are statistically indistinguishable. Conversely, a precision of *θ* = 0.05 would render that same difference highly significant. MEROS addresses this fundamental requirement by providing rigorous error estimates for occupancy and *B*-factor values, enabling researchers to distinguish genuine kinetic progressions from experimental noise. While developed with time-resolved data in mind, the utility of MEROS extends to any multi-state model where partial occupancy is a feature, such as ligand soaking, fragment screening, or multi-temperature crystallography. MEROS functions as a complementary extension to current analytical methods rather than a replacement for existing specialized tools. We can categorize its relationship to established methods into three functional groups:

Tools such as PanDDA [18] are highly effective at qualitatively identifying partial-occupancy binding events through background subtraction. MEROS could complement this by providing a downstream quantitative refinement of those identified states. Xtrapol8 [20] estimates the triggered-state occupancy and uses this to extrapolate structure factors to full occupancy. This generates clearer density maps to facilitate model building and circumvents the occupancy–*B* - Factor correlation at the coordinate-refinement stage. MEROS operates on models containing multiple simultaneous chemical states, directly refining and characterizing the occupancy of each. Furthermore, while qFit [19] excels at sampling equilibrium conformational heterogeneity (e.g., side-chain rotamers), MEROS is specifically optimized for the discrete chemical states and functional group transitions characteristic of time-resolved datasets.

Approaches like SVD-based methods and kinetic-target approaches, such as KINNTREX [16, 21], utilize neural networks to extract reaction rates directly from difference-map domains. In contrast, MEROS operates in real space on refined coordinates, producing occupancy distributions that are more intuitively suited for direct comparison with classical kinetic models.

Highly sophisticated methods like MultiXray [23] implement a Bayesian multi-state, multi-condition modeling framework to integrate multiple datasets using molecular mechanics force fields. While MultiXray offers a rigorous statistical treatment of the data-to-parameter ratio, MEROS provides a more accessible and computationally streamlined alternative for the broader community.

The primary advantage of MEROS is its simplicity and seamless integration into existing workflows. Unlike many specialized analytical methods, MEROS requires no modifications to standard refinement programs and no complex mathematical machinery beyond standard statis-tics. This modular architecture allows it to be used retrospectively. Any multi-state refinement can be re-evaluated by running the MEROS pipeline with randomized starting parameters. Despite its utility, several factors must be considered when interpreting MEROS outputs. The requirement for *N* independent refinement runs (typically *N* = 500) increases the computational workload by a factor of *N*. While this is manageable on modern multi-core workstations or compute clusters (*↘* 67 min for CTX-M-14), it may become a bottleneck for very large complexes or when seeking extreme precision (*N >* 1000). However, because the refinements are fully independent, MEROS is highly parallelizable and scales efficiently in high-performance computing environments. MEROS is not an automated state-detection tool and therefore requires the user to define the number and structural identity of the states. Consequently, its scope is currently best suited for localized regions of interest, such as active sites or hydrophobic cavities. For enzymes undergoing global structural rearrangements, or when reaction intermediates are poorly defined, the method relies on structural hypotheses. In these cases, MEROS serves as a powerful validation tool. If a hypothesized state is not supported by the data, the resulting occupancy distributions will typically fail to converge or exhibit unphysically large *B*-factors. MEROS refines a single population-averaged occupancy per state. There is an inherent risk of over-parameterization if too many independent occupancy groups are defined, particularly at low to medium resolutions. An excessive number of degrees of freedom can lead to noise fitting, where the refinement program artificially adjusts occupancies to compensate for errors elsewhere in the model. To mitigate this, users should prioritize grouping residues into chemically and physically meaningful entities to maintain a robust data-to-parameter ratio. A fundamental characteristic of MEROS is that its statistical output is conditioned on the quality of the individual refinements it orchestrates. The convergence behaviour, and therefore the width and shape of the resulting occupancy and *B*-factor distributions, are ultimately determined by the ability of the chosen refinement program to correctly minimize the crystallographic target function with respect to occupancy and *B*-factor parameters simultaneously. When the refinement program successfully resolves the occupancy–*B*-factor correlation, MEROS provides high-confidence distributions.

## Conclusion

We have presented MEROS, a general and practical framework for determining local multistate occupancies with uncertainty estimates from crystallographic data. Through systematic exploration of the joint occupancy–*B*-factor refinement landscape via Monte Carlo sampling of starting parameters, MEROS elucidates the precision with which experimental diffraction data determine each state’s population. Validation across two enzyme systems of varying complexity, the two-state ligand binding (T4 lysozyme L99A) and four-state covalent catalysis (CTX-M-14 *β*-lactamase), demonstrates that the method produces robust, reproducible occupancy distributions whose widths reflect the genuine informational content of the data. MEROS is implemented as a pipeline that wraps established refinement programs, requiring no modifications to existing crystallographic workflows. We anticipate that MEROS will be broadly useful wherever partial occupancy of enzyme states is a feature of the structural model, as in time-resolved crystallography experiments.

## Acknowledgments

All multi-temperature TR-SSX data were collected at beamline P14.2 (T-REXX) operated by EMBL Hamburg at the PETRA-III storage ring (DESY, Hamburg, Germany). We would like to thank David von Stetten for his never-tired support during data-collection. We would like to thank Pedram Mehrabi, Helen Ginn, and Nicholas Pearce for helpful discussions during preparation of MEROS.

## Author Contributions

A.P. and E.C.S. conceived and designed the MEROS method. A.P. M.S. collected diffraction data. A.P. developed the software. A.P. and M.S. performed all computations. A.P. and E.C.S. wrote the initial draft and all authors discussed and edited the manuscript.

## Declarations

The authors declare no competing financial interests.

## Data availability

The protein structures analysed in this study have been published before and are deposited in the Protein Data Bank (PDB) under the accession codes (CTX-M-14-piperacillin-3s: 20C – 9G80; 30C – 9G81; 37C – 9G82; T4L-L99A-indole-20C: 1s – 29NH; 4s – 29NJ; 40s – 29NN) [12, 29].

The source code for the MEROS algorithm is available on GitHub at https://github.com/AndiPi-1/MEROS. All other data are available from the corresponding author upon reasonable request.

