## Supplementary material for "Multi-state Ensemble Refinement for Occupancy Statistics (MEROS) in Time-Resolved X-ray Crystallography"

### Supplementary Information

#### Protein purification, expression, crystallization

##### T4 lysozyme - T4L

T4L-L99A mutant lysozyme gene was cloned in pET29b(+) vector (Genscript) and expressed in BL21 DE3 *E. coli* cells. The protein was purified by ion exchange and size exclusion chromatography with a final buffer of 0.05 M $NaH_2PO_4/Na_2HPO_4$ pH 5.5, 0.1 M NaCl, 0.002 M EDTA [32]. Batch crystallization with microseeding was performed in a total volume of 150 $\mu$L by mixing 50 $\mu$L of protein at 22 mg/mL, 45 $\mu$L of crystallization buffer consisting of 4.0 M $NaH_2PO_4/K_2HPO_4$ pH 7.0, 0.1 M *1,6*-hexanediol and 0.15 M NaCl and 5 $\mu$L of seed stock (PDB ID: 3K2R).

##### $\beta$-lactamase CTX-M-14

The production and purification of CTX-M-14 were carried out based on prior methods [33]. Briefly, E. coli BL21 (DE3) cells transformed with the pCR4:CTX-M-14 plasmid were cultured at 37 °C in LB medium containing 100 µg/mL ampicillin. Upon reaching an OD600 of 0.6-0.7, 150 µM IPTG was added to trigger protein expression, followed by a 4-hour incubation at 37 °C. Cells (5500 × g, 10 min, 4 °C) were stored at -20 °C until needed. Cells were lysed by sonication in a 20 mM MES buffer (pH 6). The cleared supernatant, after 1 h of centrifugation at 20,000 × g (4 °C), was dialysed (6–8 kDa MWCO) against a large volume of the same MES buffer over night. Cation exchange chromatography (5 mL HiTrap SP FF, Cytiva) was utilized for purification, applying a 0-50 mM NaCl gradient across 5 column volumes in 20 mM MES (pH 6). The sample was concentrated to 22 mg/mL via 10 kDa MWCO ultrafiltration (Amicon Ultra-15). Batch crystallization was performed by mixing purified CTX-M-14, crystallizing agent (40 % w/v PEG 8000, 200 mM LiSO4, 100 mM sodium acetate, pH 4.5), and undiluted seeds at a ratio of 50:45:5 (v/v/v). Homogeneous microcrystals (11-15 µm) formed in about 90 minutes. They were centrifuged (200 × g, 5 min) and resuspended in a stabilization solution (28 % w/v PEG 8000, 140 mM LiSO4, 70 mM sodium acetate, 6 mM MES pH 4.5, 15 mM NaCl) to arrest further growth. Before data collection, 0.333 M piperacillin was dissolved in initiation buffer (140 mM LiSO4, 70 mM sodium acetate, 6 mM MES pH 4.5) and kept at room temperature.

#### Ligand binding via LAMA within the 5D-SSX setup

Time-resolved serial synchrotron crystallography (TR-SSX) was performed using a microdrop nozzle for ligand droplet deposition [34] on the fixed-target HARE chips [11, 35] while the setup was in the environmental control system [12] for regulation of temperature and humidity. Briefly, microcrystals were loaded onto fixed-target HARE chips containing 20,736 features. Diffusion-based reaction initiation was driven by the 'liquid application method for time-resolved analyses' (LAMA), employing a piezo-actuated droplet injector. To ensure optimal performance, highly concentrated ligand solutions were freshly prepared, sterile-filtered, and degassed prior to application. The LAMA ejection nozzle was positioned approximately 1 mm from the chip surface and precisely aligned with the crystal wells using an on-axis infrared viewing system. Single ligand droplets (75–150 pL) were dispensed onto individual microcrystals. The automated reaction initiation sequence consisted of four steps: (1) positioning the target well, (2) triggering the detector to acquire a 5 ms pre-injection reference image ($t_0$), (3) dispensing the droplet and (4) capturing the target X-ray exposure ($t_1$) after the specified delay. The setup was within a dedicated environmental control box allowing for the regulation of temperature and humidity.

#### Data-collection and Processing

Diffraction data were collected at the PETRA-III synchrotron (DESY, Hamburg) at the EMBL P14.2 (T-REXX) endstation. Fixed-target chips were translated through a 10 × 7 µm (H × V) X-ray beam using a high-precision 3-axis SmarAct piezo stage (SmarAct, Oldenburg, Germany), and diffraction data were recorded on a Dectris Eiger 4M detector (Dectris, Baden-Daettwil, Switzerland). Diffraction data were processed using the CrystFEL (v0.10.2) [36]. Initial phases were obtained by molecular replacement in PHASER [37], using previously solved structures (6GTH for CTX-M-14; 4W51 for T4L-L99A) as search models. Subsequent structure refinement was carried out using `phenix.refine` (PHENIX v1.21.2 [25]). Iterative cycles of refinement and manual model building in `COOT` (v0.8) [38] were performed until convergence. Refinement of the occupancy was performed in `phenix.refine` by defining constrained occupancy groups for ligand-free and ligand-bound proteins, and by simultaneously refining all states in a single structure.

#### Supplementary figures

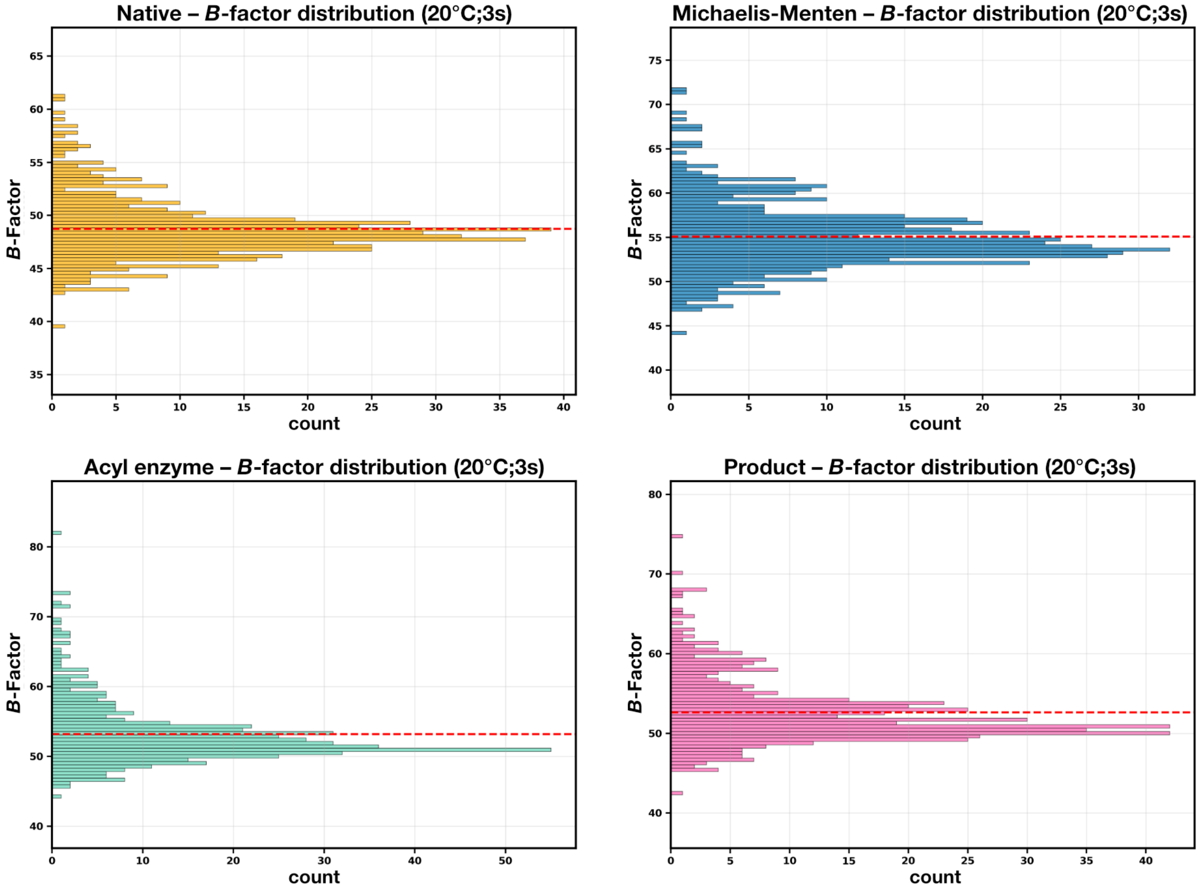


**Figure S1:** $B$-factor distribution of MEROS refinement of CTX-M-14 mixed with piperacillin at 20 °C, after a delay time of 3 seconds, with $N = 500$ refinements. The individual histograms show that the shape characteristically non-Gaussian, exhibiting a pronounced shoulder towards elevated $B$-values. The red dashed line indicates the mean value of each state.
